# Structural basis for guide RNA selection by the RESC1-RESC2 complex

**DOI:** 10.1101/2023.02.01.526499

**Authors:** Luciano G. Dolce, Yevheniia Nesterenko, Leon Walther, Félix Weis, Eva Kowalinski

## Abstract

Kinetoplastid parasites, such as trypanosomes or leishmania, rely on RNA-templated RNA editing to mature mitochondrial cryptic pre-mRNAs into functional protein-coding transcripts. Processive pan-editing of multiple editing blocks within a single transcript is dependent on the 20-subunit RNA editing substrate binding complex (RESC) that serves as a platform to orchestrate the interactions between pre-mRNA, guide RNAs (gRNAs), the catalytic RNA editing complex (RECC), and a set of RNA helicases. Due to the lack of molecular structures and biochemical studies with purified components, neither the spacio-temporal interplay of these factors nor the selection mechanism for the different RNA components is understood. Here we report the cryo-EM structure of *Trypanosoma brucei* RESC1-RESC2, a central hub module of the RESC complex. The structure reveals that RESC1 and RESC2 form an obligatory domain-swapped dimer. Although the tertiary structures of both subunits closely resemble each other, only RESC2 selectively binds 5’-triphosphate-nucleosides, a defining characteristic of gRNAs. We therefore propose RESC2 as the protective 5’-end binding site for gRNAs within the RESC complex. Overall, our structure provides a starting point for the study of the assembly and function of larger RNA-bound kinetoplast RNA editing modules and might aid in the design of anti-parasite drugs.

**Key findings:**

- The kinetoplastid mitochondrial RNA editing factors RESC1 and RESC2 resemble a group of capping enzymes that are only found in protozoans, fungi and viruses.
- RESC1 and RESC2 lack the typical catalytic residues, and only RESC2 can bind a triphosphate-nucleoside.
- We propose that the RESC1-RESC2 dimer selects guide RNAs based on their 5’-triphosphate and serves as a protective 5’-end binding complex.

## Introduction

Kinetoplastids are a clade of unicellular parasites that include *Trypanosome brucei, Trypanosoma cruzi* and *Leishmania spp*, all causative agents of neglected tropical diseases. Their distinguishing feature is the presence of the kinetoplast, a DNA-containing granule located within the large mitochondrion. Kinetoplast DNA (kDNA) comprises circular chromosomes, so-called maxicircles, which harbour protein-coding genes (1–3). Most of these genes are transcribed into cryptic pre-mRNAs that require RNA-guided RNA editing to generate fully functional transcripts for the expression of respiratory chain subunits (4–8). RNA-editing is facilitated by short guide RNAs (gRNAs) that are transcribed from small circular DNAs, termed minicircles, which are also part of kDNA. The gRNAs anneal to the pre-edited mRNA through their complementary 5’ anchor region. Sequence mismatches between pre-edited mRNA and gRNA lead to bulges that mark editing sites. These are processed by the editosome, a multi-protein machinery that generates the fully complementary, edited mRNA through the insertion or deletion of uridines (7, 9). Transcripts with multiple editing sites (“pan-editing”) will go through several rounds of editing (10). In most cases, the anchor region of the following gRNA anneals to a previously edited region, thus ensuring 3’ to 5’ processivity (8). In some cases, the editing is responsible for restoring the start codon, with alternative editing allowing for dual coding capabilities (11, 12).

The protein platform that coordinates the complex editing cycle within the editosome is the <u>R</u>NA <u>e</u>diting <u>s</u>ubstrate binding <u>c</u>omplex (RESC) (13–15). RESC consists of more than 20 different proteins and comprises three sub-modules: the guide RNA binding complex (GRBC, RESC1-7), responsible for gRNA binding and stabilisation (13, 16, 17), the RNA editing mediator complex (REMC, RESC8-13), involved in mRNA binding and processivity (18–20) and the polyadenylation mediator complex (PAMC, RESC15-18), which targets the edited mRNA to the kinetoplast polyadenylation complex (21). For the enzymatic editing reaction, the RNA editing catalytic complex (RECC) interacts with RNA-bound RESC. RECC is another large protein complex that contains multiple enzymatic activities necessary for the editing reaction: RNases for cleavage, exoUases or TUTases to delete or insert uridines, respectively, and ligases, to close the nick in the edited mRNA (Aphasizhev et al. 2003; Panigrahi et al. 2003). However, RECC alone is unable to exchange gRNAs or progress directionally on the mRNA. Pan-editing with multiple gRNAs relies for that reason on RESC (10).

The proteins RESC1 (previous nomenclature GRBC1, GAP2) and RESC2 (previous nomenclature GRBC2, GAP1) have been identified as the gRNA binding core of the RESC1-7 module (13, 22). RESC1 and RESC2 form a complex and are dependent on each other for stability. RNAi-mediated knockdown of either gene showed that both are essential for parasite growth; their depletion reduces the levels of edited mRNAs and destabilises the entire gRNA population (16, 23). Since both proteins have no sequence homology outside of the kinetoplastid family and no known domains or RNA binding motifs are apparent from their primary sequence, little is known about how they act within the RESC complex and how they select and stabilise gRNAs.

Here we report the cryo-EM structure of the *Trypanosoma brucei* RESC1-RESC2 heterodimer and show that the RESC2 subunit binds the 5’-triphosphate-nucleoside of RNA in a phosphate binding tunnel that resembles the phosphate binding site in a group of RNA capping enzymes. Since only gRNAs, but not other mitochondrial RNA species of kinetoplastids carry a 5’-triphosphate, we propose that the RESC1-RESC2 module discriminates gRNAs from the RNA pool, protects them from exonucleolytic cleavage and targets them to the editing reaction.

## Results

### The cryo-EM structure of the RESC1-RESC2 dimer

We co-expressed *Trypanosoma brucei* RESC1 and RESC2 without their mitochondrial targeting sequence in insect cells and purified the complex using a standard affinity, ion-exchange, and size exclusion chromatography protocol. The elution volume in the size exclusion suggested a dimer of RESC1 and RESC2, which we confirmed by SEC-MALS and mass photometry (Figure 1A, 1B and Supplementary Figures S1A and S1B). We solved the cryo-EM structure of the complex and built the model into the final 3.4 Å resolution map with an AlphaFold prediction of the dimeric complex as a starting point (24). The final model comprises RESC1 residues 152-175 and 185-C, and RESC2 residues 133-161 and 171-483 (Figure 1A and 1D). For each subunit, a small N-terminal domain (RESC1 46-100 and RESC2 51-107) connected by a long linker (RESC1 101-151 and RESC2 108-132) is predicted by AlphaFold but not resolved in the final cryo-EM map, suggesting flexibility within the linker region (Supplementary Figure S1C, S1D and S1E).

**Figure 1:**
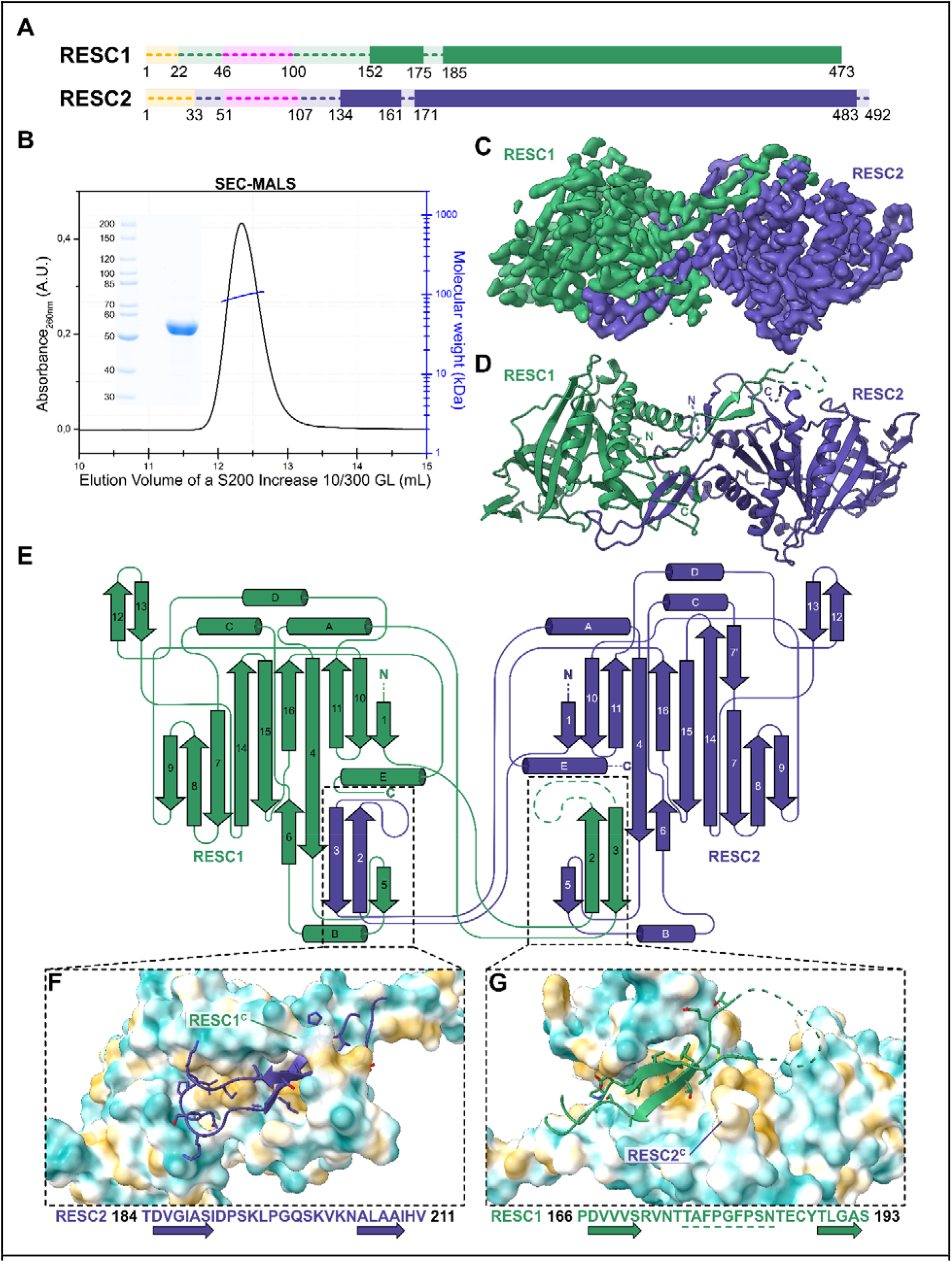
The structure of *Tb* RESC1-RESC2 heterodimer. **(A)** Linear representation of RESC1 (dark green) and RESC2 (dark blue). Regions with traced atomic models in solid colour. In yellow the mitochondrial targeting sequence, and in pink the flexibly attached N-terminal domains. Non-resolved linker regions as dashes lines. **(B)** SEC-MALS experiment with SDS-PAGE of the RESC1-RESC2 sample, indicating a molecular weight of 107 kDa, and confirming the heterodimeric oligomerisation state. **(C)** The final post-processed EM map with a 3.4 Å resolution, RESC1 is coloured in dark green and RESC2 in dark blue. **(D)** Cartoon representation of the atomic model built based on the EM map. Coloured as the map. **(E)** Structure topology diagram of RESC1-RESC2 with secondary structure elements labelled. **(F and G)** Close-up of the dimerisation interfaces, with the β-harpin of RESC2 interacting with a patch in RESC1 in F, and the β-harpin of RESC1 interacting with a cleft in RESC2 in G. Sequence of both β-harpins below, illustrating the different nature of the interaction. The unbuilt region of the RESC1 loop underlined. C-terminal regions are labelled. Surface coloured by molecular lipophilicity, with blue as hydrophilic and yellow as hydrophobic. Residue side chains coloured by heteroatom.

RESC1 and RESC2 share 35% of sequence identity (Supplementary Figure S6). In the model, the tertiary structures of both subunits resemble each other, with an RMSD of 1.159 Å. Residues important for the scaffold of the proteins are conserved throughout different kinetoplastids (Supplementary Figure S6). Each subunit comprises an imperfect β-barrel composed of 10 β-strands that are surrounded by five α-helices (Figure 1D and 1E). The inside of each barrel forms a tunnel that is accessible to solvent and possible ligands. The subunits dimerise through a domain-swapped β-harpin (two antiparallel β-sheets, β-2 and β-3, connected by a ca. 15 amino acid long loop). The dimerisation hairpins partially cover the bottom of the respective barrel. In the region of the RESC2 loop, the outer surface of the RESC1 barrel is highly negatively charged (Supplementary Figure S1K and S1L). The dimerisation interface of the subunits provides a contact area of 2426 Å^2^ and involves about 28% of the resolved domain, as calculated by PDBePISA. In each subunit, a hydrophobic patch binds the β-harpin of the other subunit, mediated by interactions between β-2 and β-3 of the hairpin with β-4, β-5 and helix B of the barrel (Figure 1E, 1F and 1G). Both β-hairpins carry hydrophobic residues for binding to the surface of the barrel, but their connection loops differ in sequence and chemical properties: the loop of RESC2 contains charged residues (K195, Q199, K201, N204) that interact with the surface of RESC1, while the loop of RESC1 contains hydrophobic residues (V173, F178, F181). The RESC2 hairpin is additionally clamped onto the surface by the C-terminus of RESC1. The RESC1 hairpin is more flexible, indicated by the lower map quality that did not allow model building for this loop (Supplementary Figure S1E and S1F). These differences in the domain swap configuration rationalise the preference for RESC1-RESC2 heterodimerisation rather than homodimerisation and the instability of the subunits when expressed alone (13, 16).

### RESC1 and RESC2 resemble RNA a specific group of RNA capping enzymes

Homology searches based on their primary sequence did not identify characterised homologues of RESC1 and RESC2, and we therefore used the PDBeFold server to find similar 3D protein structures. Best hits yielded the RNA triphosphatase domain of the yeast Cet1p and the mimivirus mRNA capping enzyme, with sequence identities of 12% or 8% and RMSDs of 2.40 or 3.14, respectively. Both enzymes are components of the mRNA 5’-capping apparatus, responsible for the hydrolysis of the 5’ triphosphate into 5’ diphosphate as the first step of the capping reaction. The proteins are members of the divalent cation-dependent family of RNA triphosphatases, defined by three conserved linear motifs, termed a, b and c. The motifs expose negatively charged residues necessary for divalent cation coordination and positively charged residues necessary for triphosphate binding to the inside of a tunnel (25–28). The barrels of RESC1 and RESC2 resemble the ones of the RNA triphosphatases, with the exception of a short insertion motif composed of α-helix D and β-strands 12 and 13 that leads to the deformation of the barrel. This structural divergence does not allow residue-by-residue superposition of RESC1 or RESC2 to the triphosphatases.

In the inner tunnel of the triphosphatases, positively and negatively charged residues are segregated in two half-cylindrical portions, while in RESC1 and RESC2 the positively and negatively charged residues are broadly distributed within the tunnel (Figure 2 A-E) (29). Furthermore, in the triphosphatases, the catalytic divalent cation is coordinated in the interior of the β-barrel by three negatively charged residues (E305, E307 and E496 in Cet1p and E37, E39 and E214 in the mimivirus enzyme) and two water molecules. A single point mutation in any of these residues inactivates the enzymes (25, 27, 28, 30). In RESC1 and RESC2, two of these three crucial residues are absent, implying that RESC1 and RESC2 are not able to coordinate divalent ions and are therefore inactive (Figure 2A-E).

**Figure 2:**
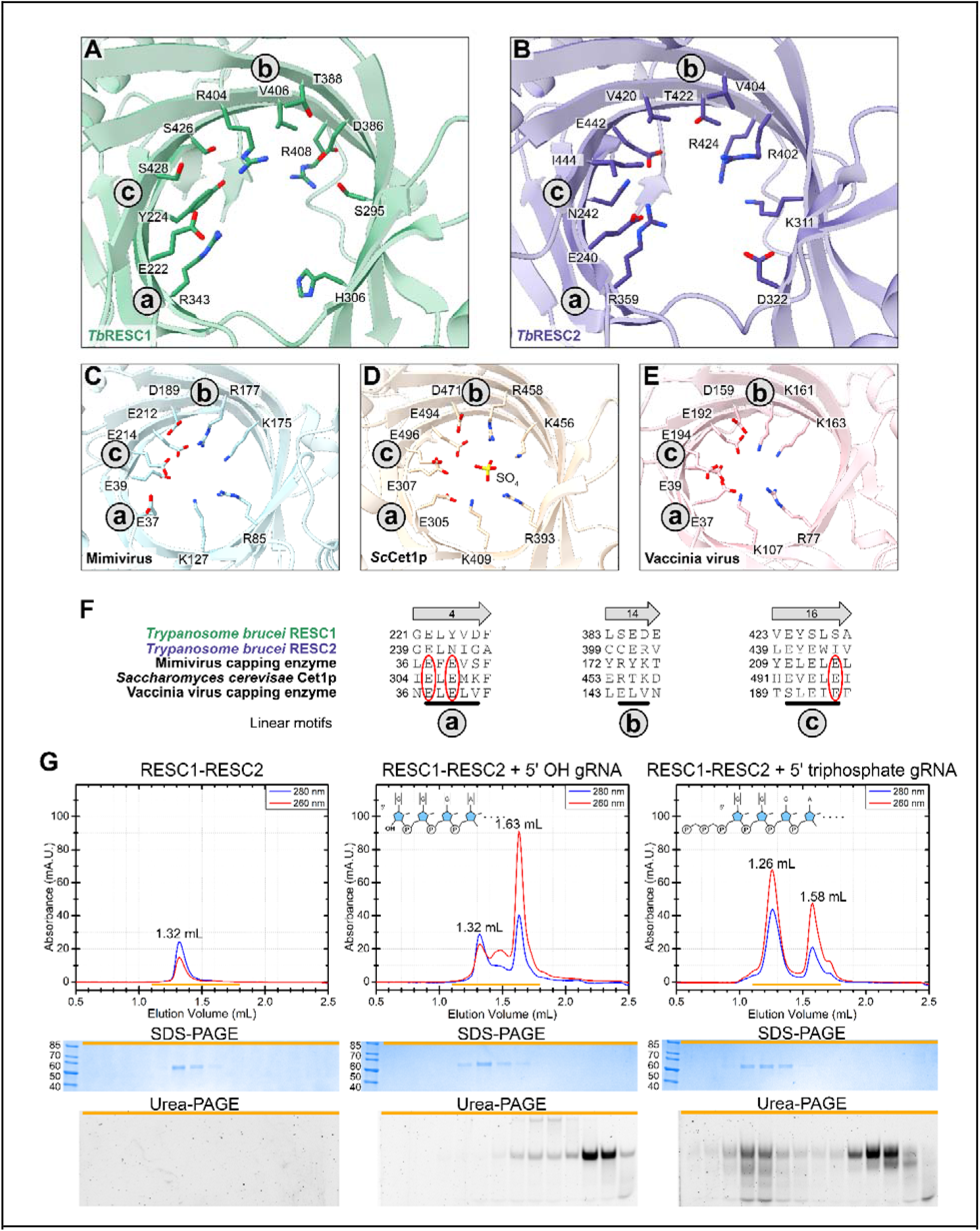
The triphosphate binding fold of *Tb* RESC1-RESC2. **(A to E)** Comparison of the chemical environment in the inner triphosphatase tunnels of **(A)** RESC1, **(B)** RESC2, **(C)** Mimivirus capping enzyme (PDB: 2QZE), **(D)** yeast Cet1p (PDB: 1D8I) and **(E)** Vaccinia virus capping enzyme (4CKC). Signature sequence motifs (called a, b and c) of the divalent cation-dependent family of RNA triphosphatases are indicated. The bound sulfate ion is shown as sticks in D. Relevant residue side chains are shown by sticks and coloured by heteroatom. **(F)** Structural sequence alignment of the three linear motifs, which are not conserved in RESC1 and RESC2. Glutamic acids were experimentally identified as fundamental for divalent cation coordination and activity in yeast Cet1p (30, 31) annotated by red circles. **(G)** Analytical size exclusion chromatography of RESC1-RESC2 with gRNA, with or without 5’ triphosphate. Yellow bars indicate the 14 fractions (from 1.1 mL to 1.8 mL) selected for SDS-PAGE for protein detection and UREA-PAGE for RNA detection. Peak shift, increase in 260:280 ratio and co-elution of the protein with the gRNA are only observed for 5’-triphosphate-containing gRNA.

### RESC1-RESC2 binds 5’-triphosphorylated RNA

Since the RESC1-RESC2 complex has been reported to bind gRNAs (13, 16), we examined our structural model for how and where the interaction might take place. In Cet1p and the mimivirus triphosphatase, the γ-phosphate of 5’ triphosphate mRNA is coordinated by four positively charged residues (R393, K409, K456, and R458 in Cet1p and R85, K127, K175, and R177 in the mimivirus enzyme) (Figure 2A-F). Since no nucleotide-bound structure of these enzymes is available, it is unclear in which direction the RNA would bind and exit from the barrel (27, 28, 30). In the RESC1 and RESC2 barrels, we observe positively charged residues in the putative phosphate-binding positions (Figure 2A-B). We therefore set out to investigate whether the RESC1-RESC2 dimer recognises RNA by binding the 5’-triphosphate. In analytical size exclusion chromatography (SEC) with *in vitro* transcribed RNA that harbours a 5’-triphosphate and a chemically synthesised RNA that harbours a 5’-hydroxyl of identical sequence, we observe complex formation exclusively with the 5’-triphosphate-containing RNA (Figure 2, Supplementary Figure S3A). We therefore conclude that the 5’-triphosphate is required for the interaction of the RESC1-RESC2 complex with RNA and that the 5’-triphosphate binding mode might be conserved between capping phosphatases and RESC1-RESC2.

### RESC2 is the subunit responsible for 5’ triphosphate binding

In a SEC-MALS experiment, the RESC1-RESC2 dimer binds 5’-triphosphorylated RNA in a 1:1 ratio, and a larger excess of gRNA did not lead to a further peak shift in a titration experiment (Supplementary Figure S3B and S3C). This suggests that only one of the two subunits coordinates gRNA. Closer inspection of the differences between the RESC1 and RESC2 structural models did not allow us to identify which subunit binds RNA, because the inner tunnel of both subunits harbours arginine and lysine residues that could mediate triphosphate binding (Figure 2A, 2B). Therefore, we reconstituted the RESC1-RESC2 complex with 5’-triphosphate RNA for cryo-EM structure determination. We treated the sample with RNase A before the final size exclusion step to remove flexible parts of the RNA that would hinder high-resolution data acquisition. A final reconstruction extended to 4.7 Å resolution. The apo-RESC1-RESC2 map was lowpass-filtered and subtracted from the gRNA-bound map. The resulting subtraction map revealed a prominent density inside the tunnel of RESC2, but not RESC1 (Figure 3A). We fitted a guanosine triphosphate (GTP) as the first nucleotide of our input sequence in the subtraction map with the γ-phosphate resembling the position of the sulfate in the *Sc*Cet1p structure (PDB: 1D8I) (32). The nucleoside moiety points towards the tunnel exit with the domain-swap hairpin. The model suggests an interaction between the nucleobase and a RESC2 loop (residues 273-279), while interaction with the sugar moiety of GTP cannot be observed. RNase A cleaves specifically before pyrimidine residues generating a 5’-triphosphate-GGG trinucleotide from our input sequence (33), but the absence of further map density suggests flexibility of the 2nd and 3rd nucleotides. Four positively charged residues of RESC2 lie in proximity to the γ-phosphate and might contribute to its coordination, as also observed in Cet1p. But two of these positions are not corresponding to positive charges in RESC1 (RESC2: K311, R359, R402, R424; corresponding RESC1: S295, R343, D386, R408), suggesting that K311 and R402, which are solely present in RESC2, might be primarily responsible for phosphate binding. To test this hypothesis, we generated the RESC2^K311S/R402D^ mutant, which mimics the RESC1 residues. Indeed, the RESC1^wt^-RESC2^K311S/R402D^ complex does not interact with 5’-triphosphate RNA (Figure 3D), marking K311 and R402 as indispensable for 5’-triphosphate coordination, and adding evidence that RESC1 is inert to phosphate binding. We conclude that RESC1-RESC2 forms a 1:1 complex with gRNA, with RESC2 as the 5’-triphosphate-interacting subunit.

**Figure 3:**
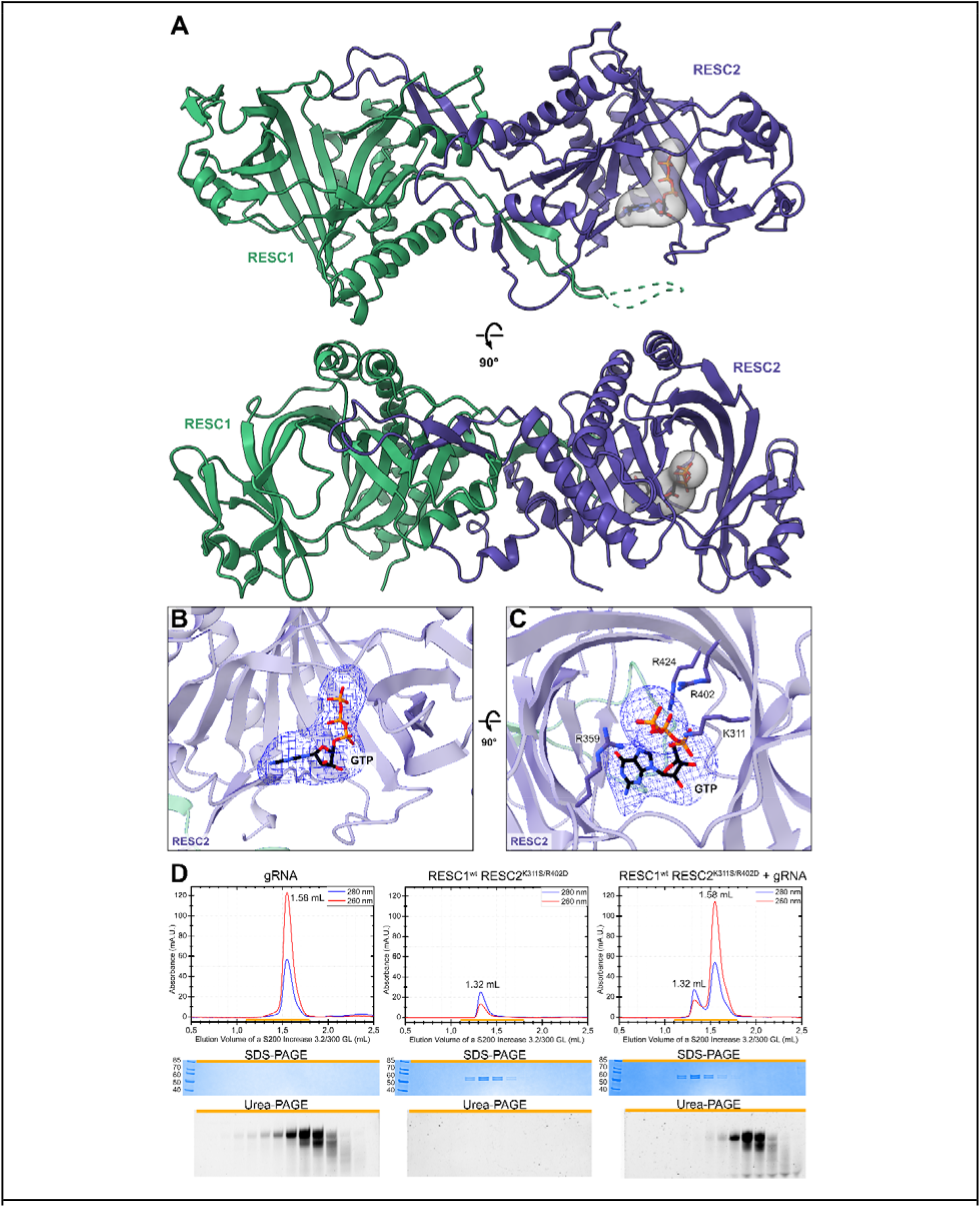
Structure of *Tb*RESC1-RESC2 bound to gRNA. **(A)** Two orthogonal views of the cartoon model of RESC1-RESC2 superimposed with the subtraction map between bound and unbound RESC1-RESC2 (in grey), with the GTP shown in sticks and coloured by heteroatom, indicating additional density exclusively inside the triphosphatase tunnel of RESC2. **(B and C)** Two orthogonal views of the triphosphatase tunnel of RESC2, with a GTP fitted in the subtraction map (blue mesh). Lysine and arginine side chains, that coordinate the γ-phosphate of the GTP, are shown in sticks and coloured by heteroatom. **(D)** Analytical size exclusion chromatography of mutant RESC1-RESC2 and gRNA, together with the SDS-PAGE for protein detection and UREA-PAGE for RNA detection. Peak shift and co-elution of the protein with the gRNA is disrupted when key RESC2 residues are mutated to the corresponding residue in RESC1 (K311S and R402D), indicating that only RESC2 is able to bind gRNA through their 5’ triphosphate.

## Discussion

### RESC1-RESC2 selects the 5’-triphosphate to recruit gRNAs to the editing process

Molecular structure studies are necessary to resolve the perplexing interplay of protein-RNA complexes in kinetoplastid mitochondrial RNA editing. Our study establishes that the 5’-triphosphate of gRNAs serves as the distinguishing feature for specific gRNA recruitment to the RNA editing machinery via binding to the RESC1-RESC2 module. In the kinetoplastid mitochondrion, gRNA is the only RNA species that is characterised by a 5’-triphosphate (9, 10, 34, 35); all other RNAs are generated through processing events that remove the transcriptionally generated 5’-triphosphate. Silencing of RESC1 or RESC2 has been shown to cause a general depletion of gRNAs (13, 16), but did not affect the editing of cytochrome oxidase subunit 2 mRNA, which employs a cis-acting guide element encoded in its 3′UTR. This finding can now be explained in light of the current study by the absence of 5’-triphosphate gRNA in the reaction (Golden and Hajduk 2005; Weng et al. 2008).

The nucleotide-bound structure of the RESC1-RESC2 complex reveals how the γ-phosphate of a 5’-triphosphate-nucleoside is coordinated in the RESC2 phosphate-binding tunnel. Future research is needed to identify additional RNA binding sites within the RESC complex or in associated factors that serve to discriminate the 5’-triphosphates of gRNAs from free phosphate or other triphosphate-nucleosides that compete at high concentrations in the cellular environment.

### RESC1-RESC2 as a gRNA 5’-end binding complex

Previous studies established that depletion of either RESC1 or RESC2 leads to a decrease of mature gRNA levels, implying that this complex stabilises gRNAs (13, 36). Accordingly, silencing of the other RESC subunits RESC4 (MRB5390), RESC5 (MRB11870) and RESC6 (MRB3010) impacts only the initial stages of pan-editing, without a consequence on gRNA abundance (13, 16, 17, 37, 38). In eukaryotic cells, mRNAs that carry a 5’-m^7^G cap are bound by cap-binding complexes for protection from exonuclease cleavage. Cap-binding complexes also act as platforms for the recruitment of other factors. In analogy, we propose that RESC1-RESC2 acts as a protective 5’-end binding complex for gRNAs, stabilising them and leading them through the RNA editing cycle.

Some RESC components seem to interact with RESC1-RESC2 in a mutually-exclusive manner. RESC6 (MRB3010), for example, has been described to be associated with RESC1-RESC2, but in co-immunoprecipitation experiments with tagged KREH2, a RESC-associated helicase, only RESC1-RESC2 and gRNA, but not RESC6 were detected (39–42). This finding might indicate a spatial-temporal resolution of different editing steps, with RESC1-RESC2 acting as one of the main hubs. This idea is supported by the detection of the RESC1-RESC2 dimer in most of the described RESC modules (21). This might be due to a permanent cap-like association with the 5’-end of the gRNA. Future structures of various larger RESC assemblies *in vitro* and *in situ* might test this idea.

Given the lack of larger RESC complex structures, we can only speculate how RESC1-RESC2 engages with other components of the RESC complex. Both RNA-dependent and RNA-independent interactions have been identified within various RESC sub-assemblies (reviewed in 9). The additional N-terminal domains and linkers of RESC1 and RESC2, which are not resolved in the structure (Supplemental figure S1C and S1D), might act to engage other proteins. The RESC1 barrel with the degenerate phosphate-binding site or its negatively charged patch on the outside could be a mediator of protein-protein interactions as well. Our study showed that the RESC1-RESC2 dimer forms a stable complex with 5’-triphosphate RNA. This assembly could serve as a starting tool to reconstitute step-by-step the editing cycle to ultimately understand this extraordinary pathway.

### The unresolved gRNA-ngRNA problem

Bidirectional transcription of the gRNA genes from minicircles generates long sense and antisense gRNA precursors with 5’ overlapping regions, which are polyuridylated by RET1 and trimmed at the 3’-end by the MPsome complex. These processing steps result in symmetric duplexes of 30-60 nucleotides containing both the sense gRNA transcript and the antisense non-guide ngRNA with uridylated 3’ tails and preserved 5’ triphosphates (35). It is currently unclear at which stage in their biogenesis gRNA and non-gRNA acquire distinguishable properties that would allow the targeted degradation of ngRNAs (21, 35). Previously a tetrameric form of RESC1-RESC2 has been observed (13, 22). Based on our structural data, the tetramer could be interpreted as two RESC1-RESC2 dimers binding to complementary RNA strands that each contain a 5’-triphosphate, potentially corresponding to gRNA-ngRNA duplexes in the cell.

### Inhibiting RESC2 phosphate binding with small molecules

RNA tunnel triphosphatases are usually only found in viruses, fungi and protozoa (25, 43). They differ substantially in terms of their structure and mechanism from other eukaryotic triphosphatases and in consequence, have been considered as potential drug targets, such as the Cet1 triphosphatase of *Trypanosoma cruzi* and *Trypanosoma bruce*i (44–46). The identified 5’-triphosphate binding residues in RESC2 appear conserved within several disease-causing kinetoplastid parasites (Supplementary Figure S6). Therefore, the RESC2 phosphate binding site could be targeted in future drug development efforts. For example, small-molecule inhibitors could be designed to compete with gRNA binding and ultimately prevent RNA editing in kinetoplastids.

## Data Availability

EM maps are available at EMDB: EMD-16592 (apo) and EMD-16593 (gRNA bound) and the apo model is deposited at PDB: 8CDP.

## Supplementary Data statement

A supplementary file with Supplementary Figures has been provided.

## Acknowledgements

We acknowledge Martin Pelosse for support in using the Eukaryotic Expression Facility at EMBL Grenoble, and Sarah Schneider for support in using the EM Facility at EMBL Grenoble. This work benefited from access to the cryo-EM platform of the Structural and Computational Biology Unit at EMBL Heidelberg. We acknowledge the European Synchrotron Radiation Facility for providing beam time on CM01 and thank Eaazhisai Kandiah for her assistance; the data from CM01 is in ESRF data portal doi.org/10.15151/ESRF-ES-884964291. This work used the platforms of the Grenoble Instruct-ERIC centre (ISBG;UAR 3518 CNRS-CEA-UGA-EMBL) within the Grenoble Partnership for Structural Biology (PSB), supported by FRISBI (ANR-10-INBS-0005-02) and GRAL, financed within the University Grenoble Alpes graduate school (Ecoles Universitaires de Recherche) CBH-EUR-GS (ANR-17-EURE-0003). We thank Caroline Mas for access to the Biophysical platform. We thank Aline Le Roy and Christine Ebel, for assistance and access to the Protein Analysis On Line (PAOL) platform. We thank Life Science Editors for editing services (www.lifescienceeditors.com). The work was supported by a grant from the French Agence Nationale de la Recherche to E.K. (ANR-20-CE11-0016). The authors thank the Kowalinski lab members for discussions and comments throughout the project.

## Material and Methods

### Protein Expression and Purification

RESC1 (Tb927.7.2570) and RESC2 (Tb927.2.3800), without their mitochondrial targeting sequences (RESC1 22-473, RESC2 33-492), and all mutants thereof, were co-expressed in Hi5 cells using the Bigbac insect cell expression system (47, 48). Cells were resuspended in lysis buffer (20 mM HEPES pH 7.5; 200 mM NaCl; 2 mM MgCl_2_; 20 mM Imidazole; 2% Glycerol; supplemented with 1 mM of PMSF, 2 µg/mL of DNase and 25 µg/mL of RNase) and lysed by sonication (5 minutes, 30% Amplitude, 5 seconds on, 15 seconds off; Vibra-cells, Sonics). The lysate was clarified by centrifugation (40,000 g, 30 minutes) and loaded into a 5 ml HisTrap HP affinity chromatography column (Cytiva), that was washed with 5 column volumes of lysis buffer, 10 volumes of high salt buffer (20 mM HEPES pH 7.5; 1 M KCl) and 5 column volumes of low salt buffer (20 mM HEPES pH 7.5; 200 mM NaCl; 2 mM MgCl_2_; 50 mM Imidazole). The sample was eluted with an imidazole gradient and dialysed to a low salt ion exchange buffer (20 mM HEPES pH 7.5, 100 mM NaCl, 2 mM MgCl_2_, 1 mM DTT), when needed, his-tag was cleaved during dialysis with 3C protease. After dialysis, the sample was injected in a HiTrap Q HP anion exchange chromatography column (Cytiva), and eluted with a NaCl gradient. The protein fractions were pooled, aliquoted, flash-frozen in liquid nitrogen and stored at −80°C. As a final purification step, frozen aliquots were thawed and injected in a Superdex 200 size exclusion column (Cytiva), pre-equilibrated with the desired buffer for downstream applications (20 mM HEPES pH 7.5, 2 mM MgCl_2_, 1 mM DTT and different concentrations of NaCl). All steps were performed at 4°C.

### SEC-MALS

The size exclusion chromatography - multiple angle light scattering (SEC-MALS) experiments were performed with a Superdex 200 Increase 10/300 GL column, in SEC buffer (20 mM HEPES pH 7.5, 50 mM NaCl, 2 mM MgCl_2_, 1 mM DTT), coupled to MALS detector miniDAWN™ TREOS (WYATT). The sample (2 mg/mL, 50 µL) was injected into the column with a 0,5 mL/min flow. The data were processed using the ASTRA V software (Wyatt Technology). For the complex of RESC1-RESC2 and gRNA, RESC1-RESC2 at 15 µM and gRNA at 60 µM. The mass was calculated by two-component analysis.

### Analytical SEC

Analytical size exclusion chromatography to analyse RNA binding was performed in a Superdex 200 increase 3.2/300 GL size exclusion column (Cytiva) connected to an AKTA Micro system (Cytiva) and pre-equilibrated with SEC buffer (20 mM HEPES pH 7.5, 50 mM NaCl, 2 mM MgCl_2_, 1 mM DTT). Exclusively for the experiments with RESC1^wt^/RESC2^K311S^, the buffer contained 400 mM of NaCl. Different combinations of RESC1-RESC2 and gRNA (gND7[506] (49)), at final concentrations of 10 µM and 12,5 µM, 10 µL) were injected in the column with a 0,05 mL/min flow, and the elution was fractionated in 50 µL fractions. Relevant fractions were collected and analysed by SDS-PAGE and coloured with coomassie blue for protein content or analysed by UREA-PAGE, coloured with SYBR Gold.

### Mass photometry

The Mass photometry experiment was performed in the Refeyn mass photometry equipment, with RESC1-RESC2 in a final concentration of 100 nM. The contrast-to-mass conversion was achieved by calibration using a native protein ladder, following the manufacturer’s guidelines.

### In vitro transcription of gRNAs

The gRNAs used in this work were either synthesised by Biomers (synthetic gRNA - without 5’ triphosphate) or in-vitro transcribed using the MEGAshortscript™ T7 Transcription Kit (Invitrogen) (transcribed gRNA - with 5’ triphosphate). Oligonucleotides containing the sequence of the gRNA gND7(506) (GGGAUAUAAAUACGAUGUAAAUAACCUGUAGUA-UAGUUAGUGUAUAUAGUGAAA) and a T7 promoter were annealed to make partially single-stranded templates. Following the manufacturer guidelines, the templates were added to the reaction to a final concentration of 200 nM, followed by nuclease-free water, nucleotide solutions, reaction buffer and T7 Enzyme mix, and incubated at 37°C for 4 hours. After transcription, the reaction was mixed with a gel loading buffer and loaded in a 7M Urea-PAGE. The band of RNA was visualised in a TCL place using the U.V. shadowing method, cut from the gel and eluted in gel elution buffer (0.5 M ammonium acetate, 1 mM EDTA, 0.2% SDS) at 37°C, overnight. After elution, the RNA was precipitated with isopropanol and resuspended in nuclease-free water.

### Cryo-EM sample preparation

For Cryo-EM sample preparation, RESC1-RESC2 heterodimer (2 mg/mL, 50 µL) was injected in a Superdex 200 Increase 3.2/300 size exclusion column (Cytiva) pre-equilibrated with SEC buffer (20 mM Hepes pH 7.5; 50 mM NaCl; 2 mM MgCl2; 0.5 mM TCEP). UltrAuFoil R 1.2/1.3 300 Au mesh (Quantifoil) grids were glow discharged with residual air for 30 seconds, on each side in a PELCO easiGlow device operated at 30 mA. 2 µL of the sample were applied in each side of the grid before blotting for 3 seconds (blot force 0) and vitrified in liquid ethane using a Vitrobot MARK IV (Thermo Fisher Scientific) operated at 4 °C and 100% humidity.

For the gRNA bound RESC1-RESC2 sample preparation, RESC1-RESC2 heterodimer (2 mg/mL) was incubated with in-vitro transcribed gRNA in a 1:1,2 protein : RNA molar ratio, for 30 minutes, 4 °C. After incubation, 1 µL of RNase A (Invitrogen) was added, and the sample was incubated for another 30 minutes before being injected in a Superdex 200 Increase 3.2/300 size exclusion column (Cytiva) pre-equilibrated with SEC buffer (20 mM Hepes pH 7.5; 50 mM NaCl; 2 mM MgCl2; 0.5 mM TCEP). Sample vitrification was performed the same way as for the apo sample.

### Cryo-EM Data acquisition

For the apo RESC1-RESC2 complex, micrographs were collected with a 300kV Titan Krios (Thermo Fisher Scientific) electron microscope at EMBL Heidelberg, equipped with a K2 Summit direct electron detector and a GIF Quantum energy filter (Gatan). Data were acquired using serialEM (50) at a nominal magnification of 165,000, resulting in a pixel size of 0.81□Å. For a better sampling of particle orientations, the stage was tilted 30°; to reduce beam-induced motion, the beam size was increased to 650 nm. Movies were acquired for 12 seconds at a dose of 3 e/pixel/s, resulting in a total dose of 63.00 e/A^2^ at the sample level, fractionated into 100 movie frames. Three movies were acquired per hole, for a total of 6,145 movies, with a defocus range from −1.2 to −2.5 µm.

For the gRNA bound RESC1-RESC2, micrographs were collected with a 300kV Titan Krios (Thermo Fisher Scientific) electron microscope at ESRF Grenoble, equipped with a K3 direct electron detector and a Quantum-LS energy filter (Gatan). Data were acquired using EPU at a nominal magnification of 105,000, resulting in a pixel size of 0.84□Å (51). For a better sampling of particle orientations, the stage was again tilted to 30°; to reduce beam-induced motion, the beam size was increased to 1100 nm. Movies were acquired for 2.3 seconds at a dose of 19.3 e/pixel/s, resulting in a total dose of 62.90 e/A^2^ at the sample level, fractionated into 60 movie frames. One movie was acquired per hole, for a total of 4,420 movies, with a defocus range from −1.2 to −3.5 µm.

### Cryo-EM Data processing

For the unbound RESC1-RESC2 data, movies were motion corrected, and CTF was estimated in Relion 3.1 (52). A total of 2,109,828 particles were picked using the BoxNet2Mask_21080918 model (53) and extracted and binned (300-pixel box, binned to 100-pixel box) using Relion. Extracted particles were imported to cryoSPARC (54) and used to generate 5 ab-initio reconstructions. Particles from 4 of the 5 classes were classified by a hetero refinement job, from which a class with broken particles were discarded. The remaining 1,236,397 particles were used to generate 5 ab-initio reconstructions followed by hetero refinement from which the particles of 2 classes, for a total of 631,107 particles, were selected. Selected particles were unbinned by re-extraction in Relion, imported to cryoSPARC, and 3D classified using ab-initio reconstruction into 4 classes. The 2 highest resolution classes were once more 3D classified using ab-initio reconstruction into 4 classes. The 2 highest resolution classes, which resulted in very similar reconstructions, were pooled together for a total of 447,858 particles, and used in a non-uniform refinement, followed by a local refinement. The particles, masks and maps were imported to Relion for 3 rounds of CTF refinement and Bayesian polishing (55), resulting in a final refined map with an estimated resolution of 3.42 A as calculated by Relion gold standard FSC method.

For the gRNA bond RESC1-RESC2, movies were motion corrected, and the CTF was estimated in Relion 3.1 (52). A total of 1,514,680 particles were picked using the BoxNet2Mask_21080918 model (53) and extracted and binned (300-pixel box, binned to 100-pixel box) using Relion. Extracted particles were imported to cryoSPARC (54) and used to generate 5 ab-initio reconstructions, which were further classified by a hetero refinement job. This particle sorting process was repeated another 3 times, and the remaining 497,550 particles were imported to Relion and unbinned. Unbinned particles were classified one last time in cryoSPARC, where it was possible to see a density for the gRNA in 2 of the 5 classes, adding to 202,502 particles. These particles, masks and maps were imported to Relion for 4 rounds of CTF refinement and Bayesian polishing (55), resulting in a final refined map with an estimated resolution of 4.7 A as calculated by Relion gold standard FSC method.

### Model building

The final map was sharpened with Relion, blurred with Coot 0.9.5 (56), or post-processed with DeepEMhancer (57) for easier interpretation. For the model building, an Alphafold model of RESC1-RESC2 heterodimer served as a starting point. Regions with low predicted IDDT were deleted, and the model was refined against the non-sharpened map in an interactive manner with Ramachandran and secondary structure restraints, using both Coot 0.9.5 and Phenix (58). RMSD between input alphafold model and final model is 0.931.

For the gRNA bound RESC1-RESC2, the apo structure was rigid-body fitted in the final map, and the 5’ triphosphate, sugar and first base of the gRNA were refined inside the observed extra density. For visualisation, a subtraction map was generated by lowpass filtering the apo RESC1-RESC2 followed by volume subtraction and removal of noise induced by the difference in resolution, in ChimeraX (59).

Alignments were made with UCSF Chimera or clustalw, and displayed with ESPript (60–62).

## Author contributions

EK designed the study. EK, LGD, YN and LW conducted experimental work. FW collected the tilted cryo-EM data at EMBL. LGD processed the cryo-EM data and built the model. LGD and EK interpreted the data and wrote the manuscript.

