## Supplementar Information for "Structural basis for guide RNA selection by the RESC1-RESC2 complex"

**Supplementary figure S1: Biochemical and structural characterisation of RESC1-RESC2 heterodimer. (A)** Refeyn mass photometry measurement of RESC1-RESC2 molecular weight in solution, indicating an approximately 100 kDa particle. **(B)** Example of size exclusion chromatography used as a final step of the purification of RESC1-RESC2, from a S200 16/600 GL column. **(C)** Cartoon representation of the Alphafold prediction of RESC1-RESC2, coloured by lDDT, from 0.2 (blue) to 0.98 (red), with corresponding PAE plot. **(D)** Alphafold prediction fitted in the deepEMhancer post-process EM map by ChimeraX, with no density for the flexibly attached N-terminal domains. **(E and F)** Detail of the dimerisation interfaces β-harpin in RESC1 and RESC2, with the deepEMhancer post-process EM map around the β-harpin represented as blue mesh. The lower quality of the EM map in the region of RESC1's hairpin suggests the loop to be more flexible than in RESC2. **(G, H and I)** Example of the deepEMhancer post-process EM map, as a blue mesh, of a helix and two strands, with sticks coloured by heteroatom. **(J)** Dimerisation interface in RESC1, with RESC1 represented as surface and coloured by charges, and RESC2 as cartoon. **(K)** Dimerisation interface in RESC2, with RESC2 represented as surface and coloured by charges, and RESC1 as cartoon. **(L)** 4 views of the RESC1-RESC2 model fit in the final deepEMhancer post-process EM map.

**Supplementary figure S2: Processing scheme of the cryo-EM structure determination of RESC1-RESC2. (A)** Example of a representative micrograph, resulting from the averaging of the motion-corrected frames of a movie. **(B)** Example of 2D classes made with particles picked by warp. **(C)** Particle sorting process by ab-initio and heterogeneous refinements. **(D)** Polishing steps of the final particle stack, followed by post-processing in Relion or with DeepEMhancer. **(E)** Local resolution map, as calculated by Relion and displayed in ChimeraX. **(F)** Angular distribution map of particle orientation of the final refinement step in Relion. Map recalculated in cryoSPARC for representation proposes. **(G)** Final resolution of the reconstruction, as calculated by Relion Gold Standard Fourier Shell Correlation, with a resolution of 3.4 Å when adopting a cutoff of 0.143.

**Supplementary figure S3: Biochemical characterisation of RESC1-RESC2 5' triphosphate binding. (A)** Analytical size exclusion chromatography of RESC1-RESC2 with gRNA, with or without 5' triphosphate. Control experiments for Figure 2G are included. Chromatograms, full images of the SDS and Urea PAGEs. **(B)** Size exclusion titration of different molar ratios of gRNA on RESC1-RESC2. No further peak shift is detected above 1:1 gRNA:protein ratio. **(C)** SEC-MALS of the RESC1-RESC2 complex with gRNA in a 1:4 gRNA:protein ratio. The calculated molar mass of the particles were 18.6 ± 0.2 kDa, 102 ± 1 kDa and 113 ± 1 kDa for the gRNA, RESC1-RESC2 and their complex, respectively.

**Supplementary figure S4: Processing scheme of the cryo-EM structure determination of RESC1-RESC2 bound to 5' triphosphate gRNA. (A)** Example of a representative micrograph, resulting from the averaging of the motion-corrected frames of a movie. **(B)** Example of 2D classes, particles picked by warp. **(C)** Particle sorting process by ab-initio and heterogeneous refinement. **(D)** Polishing steps of the final particle stack, followed by post-processing, in Relion or with DeepEMhancer. **(E)** The local resolution map, as calculated by Relion and displayed in ChimeraX, indicates that the triphosphate-bound tunnel has one of the highest resolutions of the reconstruction. **(F)** Angular distribution map of particle orientation of the final refinement step done in Relion. Map recalculated in cryoSPARC for representation proposes. **(G)** Final resolution of the reconstruction, as calculated by Relion Gold Standard Fourier Shell Correlation, with a resolution of 4.7 A when adopting a cutoff of 0.143.

**Supplementary figure S5: Biochemical characterisation of RESC2 5' triphosphate binding.** **(A)** Analytical size exclusion chromatograms of $RESC1^{wt}$-$RESC2^{K311S/R402D}$ mutant complex, complementing the Figure 3D, with the controls, and the full image of the SDS and Urea PAGE. **(B)** Analytical size exclusion chromatograms of $RESC1^{wt}$-$RESC2^{K311S}$ complex. gRNA binding in this single mutant was affected but not abolished. This experiment was performed in a buffer containing 400 mM NaCl.

**Supplementary figure S6: Sequence alignment of RESC1 and RESC2 from different kinetoplastida members.** Start residue of the RESC1 and RESC2 constructs annotated as yellow diamonds, start residue of cryo-EM based model annotated by green and blue triangle for RESC1 and RESC2 respectively. Flexible attached N-terminal domains are indicated in pink. Dimerization β-harpin coloured as green and blue, for RESC1 and RESC2, respectively. Signature motifs (a, b and c) of the divalent cation-dependent family of RNA triphosphatases are indicated. The four observed 5'- triphosphate binding residues in RESC2 are indicated with a red star, the $RESC2^{K311S/R402D}$ mutant residues as a red star inside a circle. Red boxes indicate 100% conservation of RESC1 and RESC2 in all shown species.

### Supplementary Figure S1

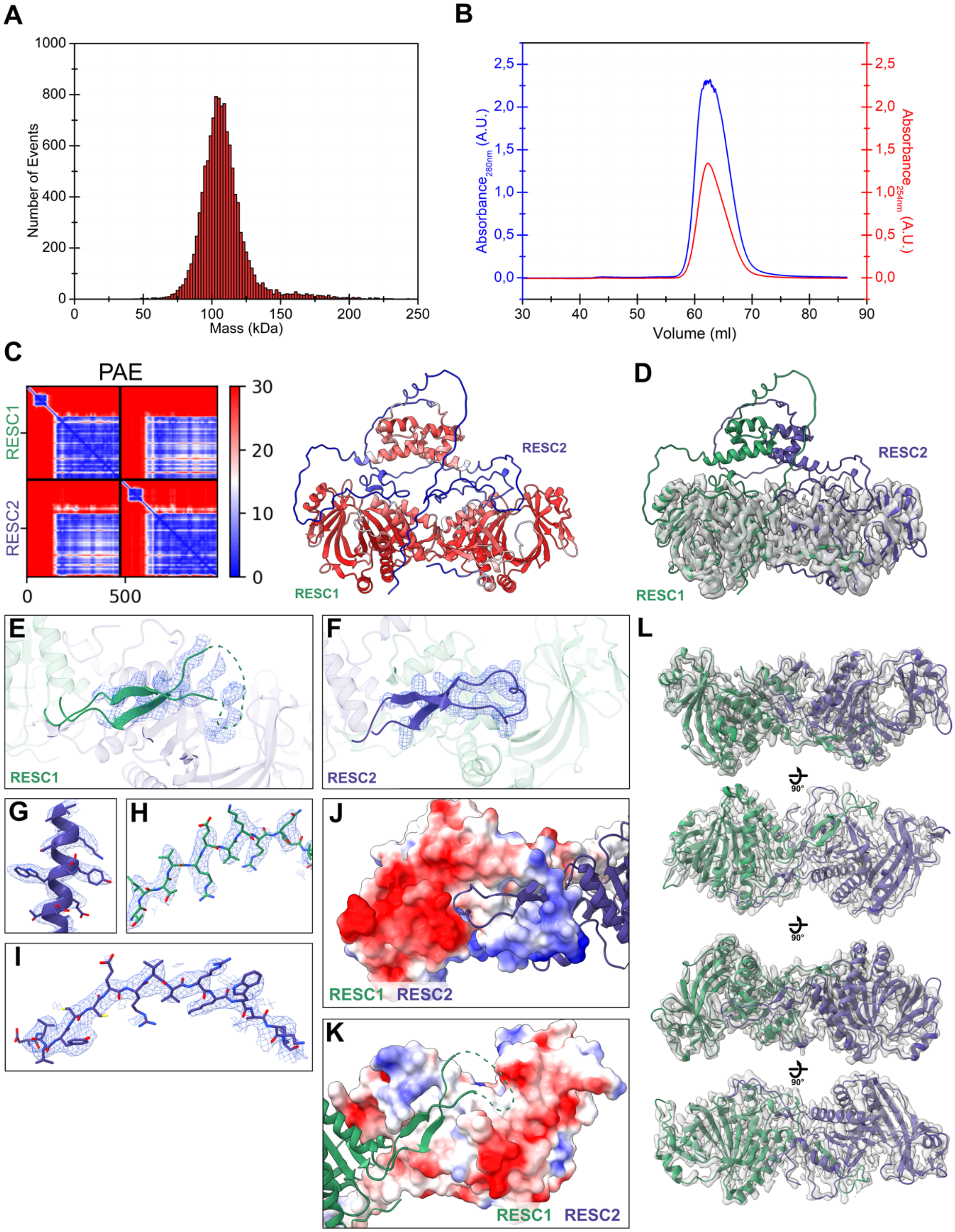

### Supplementary Figure S2

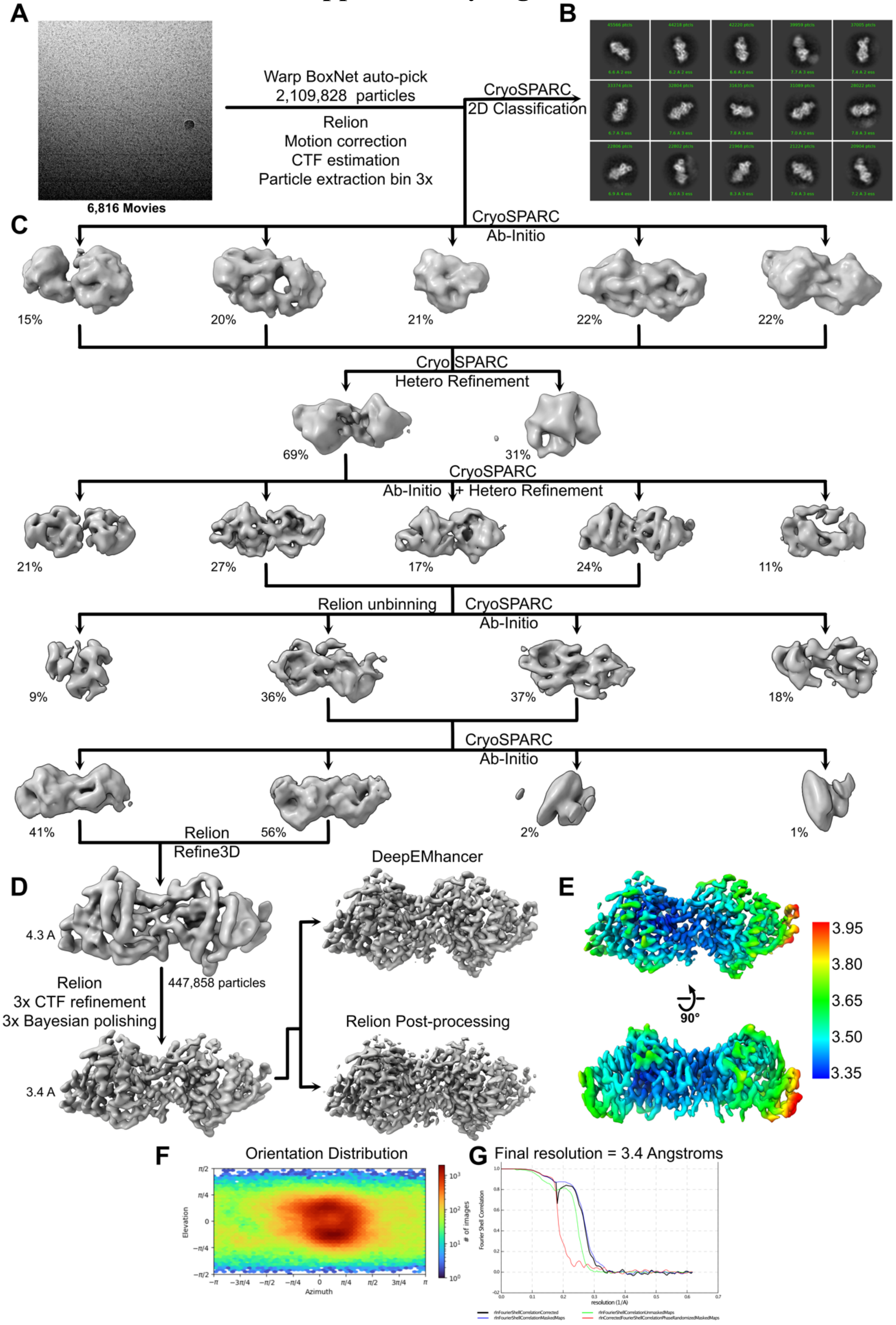

### Supplementary Figure S3

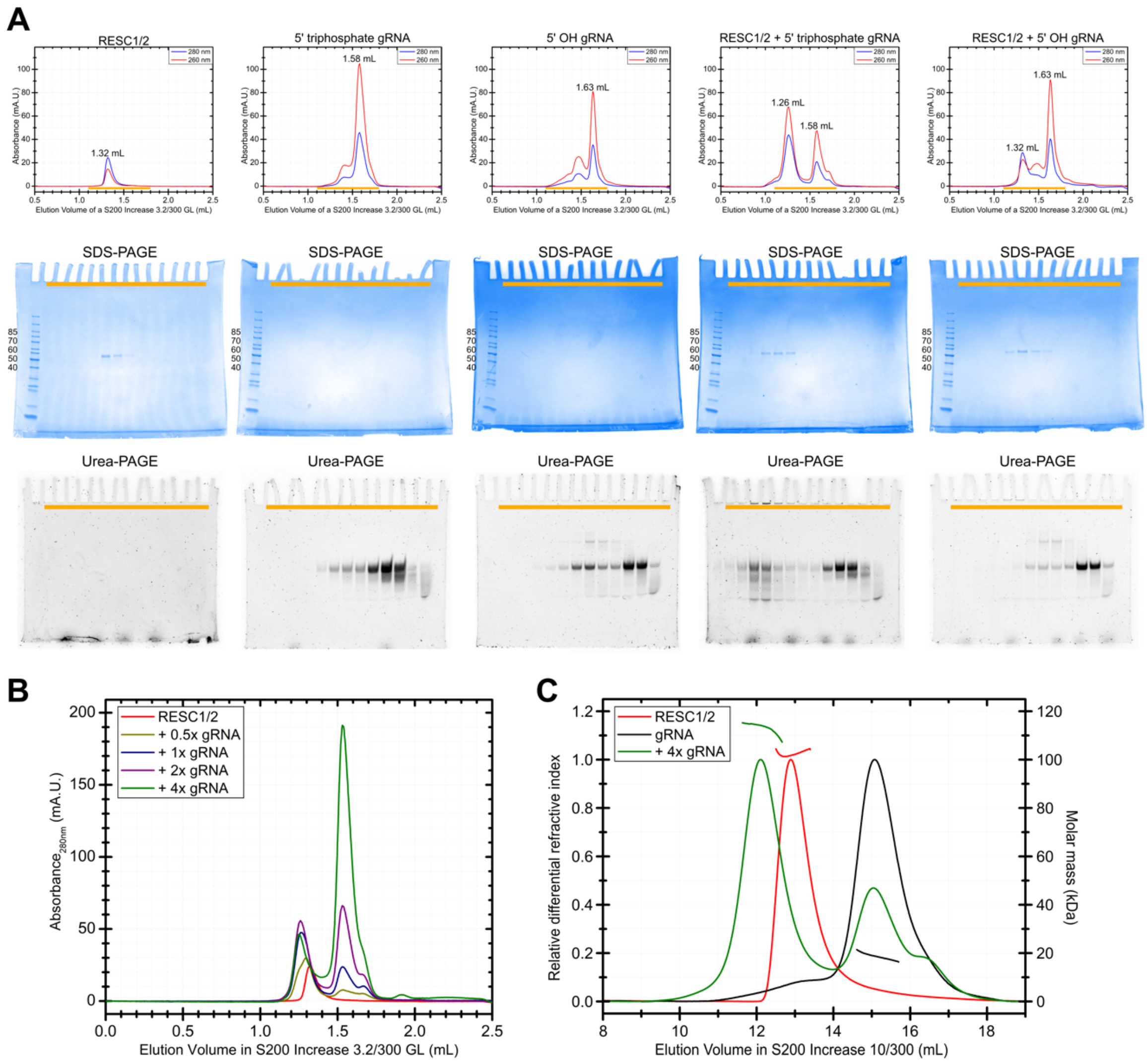

### Supplementary Figure S4

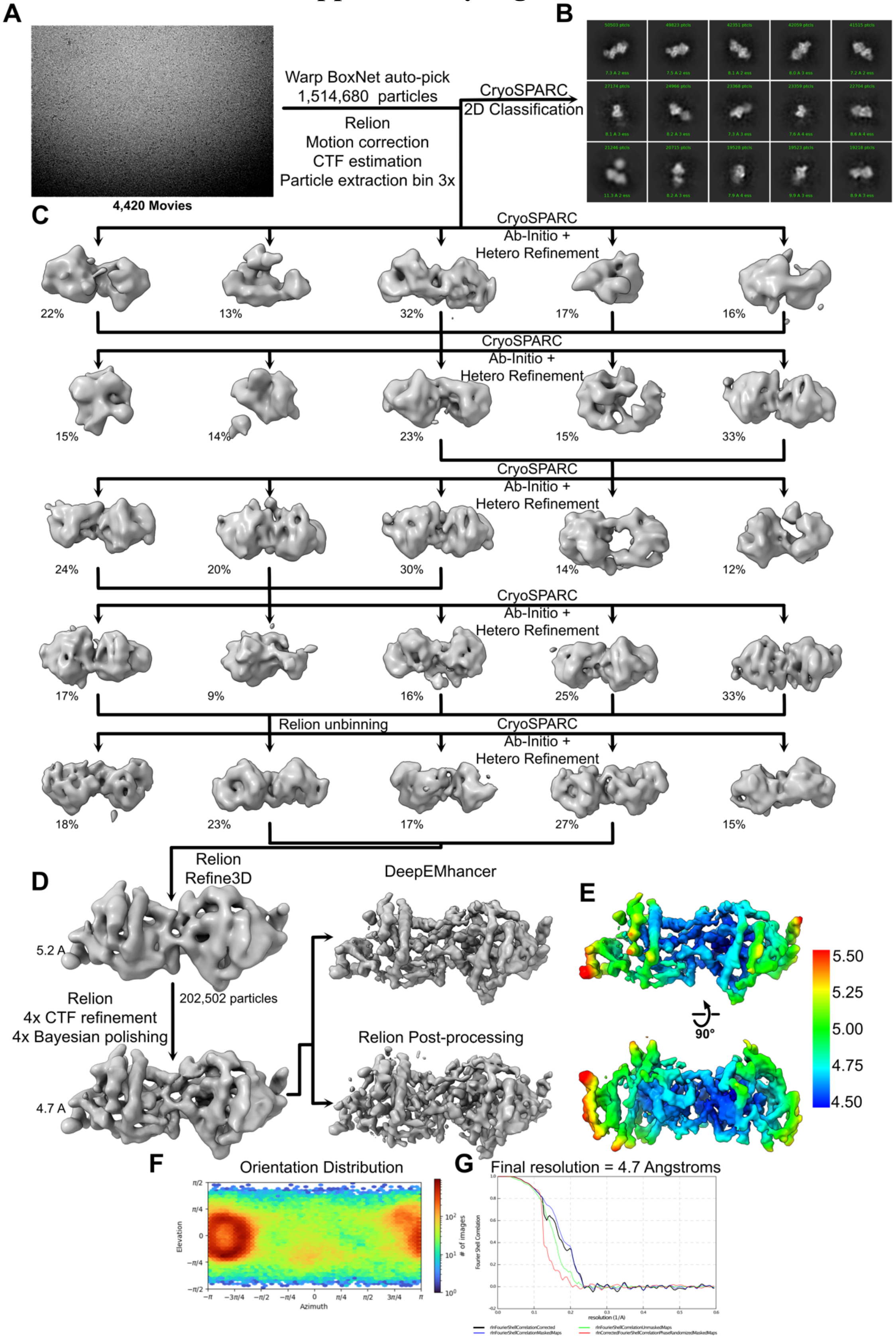

### Supplementary Figure S5

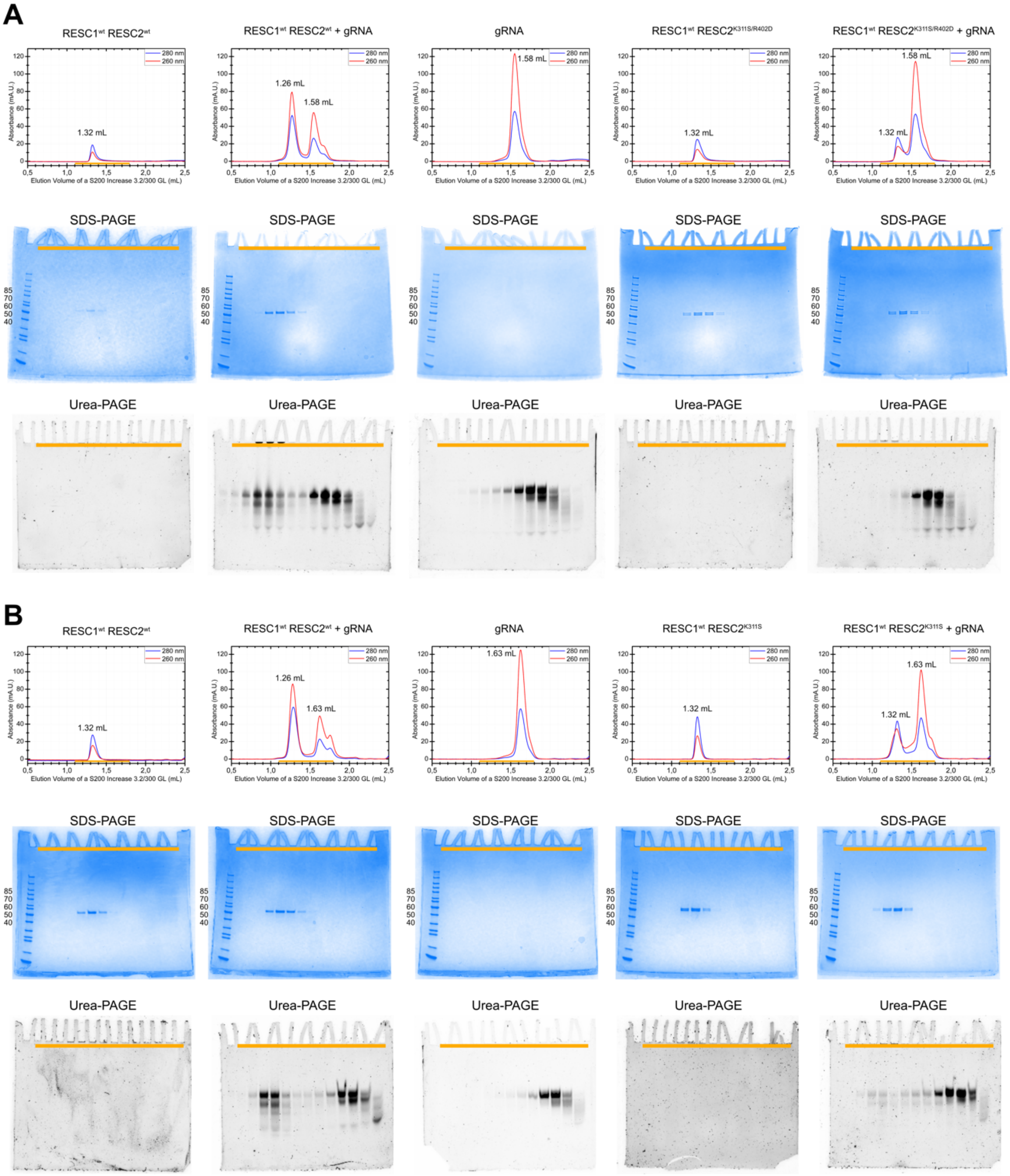

### Supplementary Figure S6

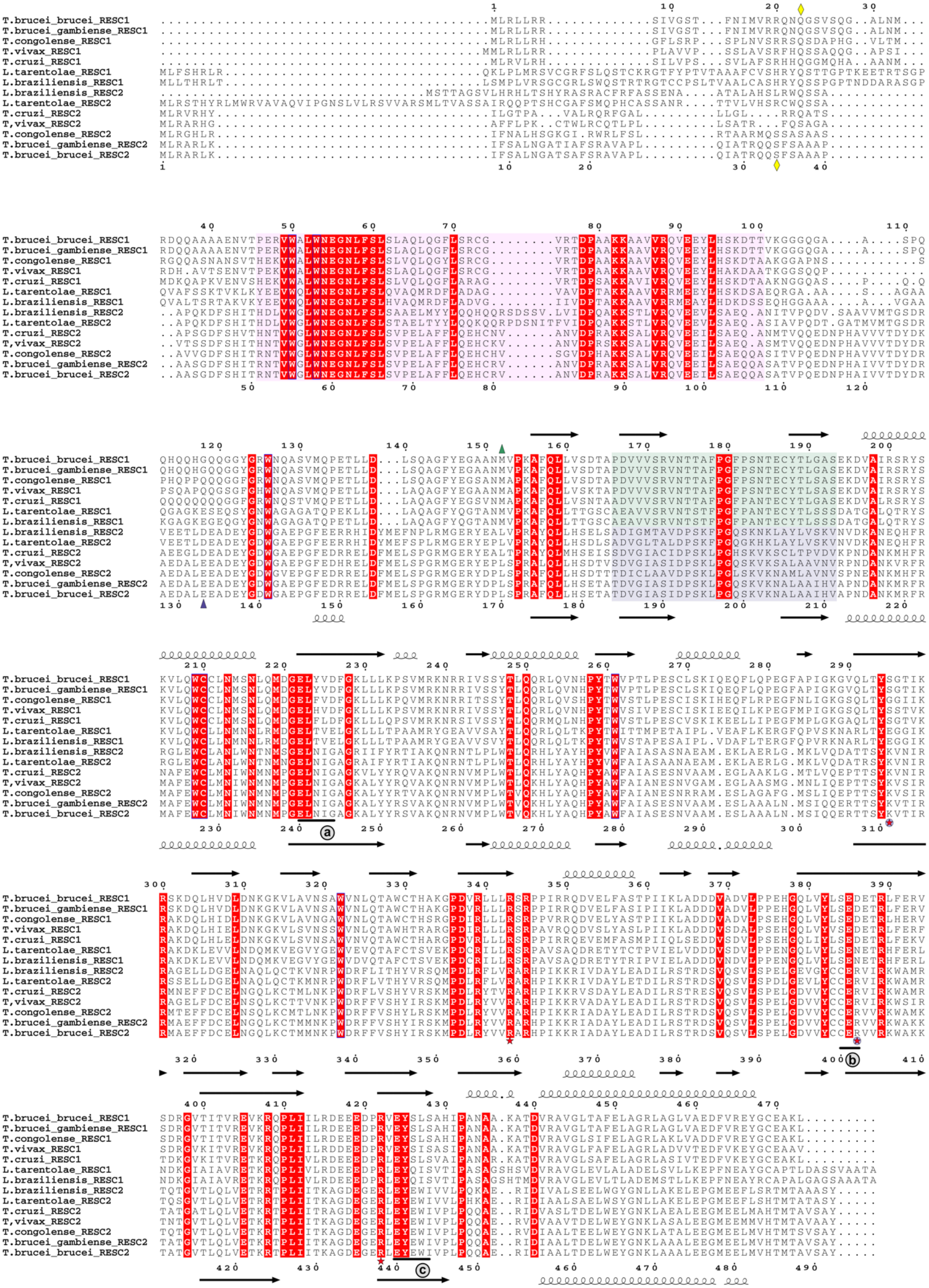

**Supplementary Table 1. Summary of Cryo-electron microscopy data collection and refinement statistics**

| **Data Collection** | **RESC1-RESC2** | **RESC1-RESC2 + gRNA** |
|---|---|---|
| Microscope | Titan Krios | Titan Krios |
| Voltage | 300 kV | 300 kV |
| Camera | Quantum-K2 | K3 |
| Energy filter | GIF | Quantum-LS |
| Spot size | 8 | 8 |
| Beam size | 650 nm | 1100 nm |
| Pixel size (A/pix) | 0.81 | 0.84 |
| Preset defocus range | -1.2 to -2.5 | -1.2 to -3.5 |
| Stage tilt | 30 deg | 30 deg |
| Dose rate (e-/pixel.s) / total exposure (e-/A2) | 3 / 63 | 19.3 / 62.9 |
| Number of frames per movie | 100 | 60 |
| Number of movies | 6,145 | 4,420 |
| **Data Processing** | **RESC1-RESC2** | **RESC1-RESC2 + gRNA** |
| Initially warp autopick particles | 2,109,828 | 1,514,680 |
| Final number of particles | 447,858 | 202,502 |
| Resolution FSC | 3.4 | 4.7 |
| Sharpening method | DeepEMhancer | DeepEMhancer |
| EMDB accession number | EMD-16592 | EMD-16593 |
| **Refinement** | **RESC1-RESC2** | |
| PDB accession number | 8CDP | |
| No atoms | 5200 | |
| Residues (protein) | 654 | |
| CCbox, CCmask, CCvolume | 0.89, 0.83, 0.83 | |
| R.M.S.D. | | |
| Bond lengths | 0.003 | |
| Bond angles | 0.620 | |
| Ramachandran favoured (%) | 97.52 | |
| Ramachandran allowed (%) | 2.48 | |
| Ramachandran outlier (%) | 0 | |
| MolProbity score | 1.69 | |
| Clash score | 12.00 | |
